# The β-lactam Ticarcillin *is a Staphylococcus aureus* UDP-N-acetylglucosamine 2-epimerase binder

**DOI:** 10.1101/2021.11.05.467346

**Authors:** Erika Chang de Azevedo, Alessandro S. Nascimento

## Abstract

Infectious diseases account for 25% of the causes of death worldwide and this rate is expected to increase due to antibiotic resistance. Among the bacteria associated with healthcare infections, *Staphylococcus aureus* is a prevalent pathogen and about 50% of the isolates are found to be methicillin-resistant. Here we describe the identification of ticarcillin as a weak binder of the *S. aureus* UDP-N-acetylglucosamine 2-epimerase. After a docking screening, ticarcillin was identified as a ligand in using the recently proposed isothermal analysis of differential scanning fluorimetry data. Finally, an equilibrium MD simulation confirmed the docking binding mode as a stable pose, with large contributions to the binding energy coming from interactions between Arg206 and Arg207 and the carboxylate groups in ticarcillin.

## 1. Introduction

The Gram-positive bacteria *Staphylococcus aureus* is listed by the Center for Disease Control and Prevention (CDC) in the USA among those bacteria that represent a serious threat to humans due to their high rates of observed antibiotic resistance [1]. The number of infections caused by methicillin-resistant *S. aureus* (MRSA) strains per year reaches 80,000, resulting in about 11,000 deaths [1]. Also, according to the survey taken by the National Healthcare Safety Network at the Centers for Disease Control and Prevention, USA, *S. aureus* was ranked as the second most frequent organism causing healthcare-associated infection, after *Escherichia coli*, and was responsible for about 12% of the reported infections [2]. Also worth of note, about 50% of the S. aureus isolates in diverse infection sites were found to be resistant to methicillin [2].

The estimated number of people with important infections caused by antibiotic-resistant bacteria reached 2,000,000 in 2013, with 23,000 deaths directly caused by these infections [1]. In Europe, about 25,000 deaths were related to antibiotic-resistant bacteria in 2007 by the European Center for Disease Prevention and Control [3]. The estimated number of deaths per year caused by antibiotic resistance globally reaches 700,000, highlighting the burden caused by resistant bacteria [4]. In addition, Thorpe and coworkers estimated that antibiotic resistance results in an additional $1,383 to the cost of the treatment per patient, resulting in an annual cost of 2.2 billion dollars in the US [5].

The bacterial cell envelope, the first line of defense for bacteria against changes in the environment, is also the target of the first-choice antibiotic for treating many *S. aureus* infections. The beta-lactam antibiotics act by binding and inhibiting the penicillin-binding proteins (PBPs) [6], impairing the transpeptidase activity and resulting in the inhibition of the peptidoglycan synthesis. The cell envelope in Gram-positive bacteria such as *S. aureus* is also decorated with the wall teichoic acid (WTA) [7], which are polymers containing 40-60 polyol repeats and can account for up to 60% of the cell wall mass [7]. The association between beta-lactam antibiotic resistance and the composition of the WTA has been shown, although the precise mechanisms underlying this association are not completely clear yet. Brown and coworkers showed that the deletion of the glycosyltransferase (GT) TarS, responsible for the decoration of the WTA with β-O-GlcNAc, can re-sensitize MRSA strains to beta-lactam antibiotics [8]. Additionally, it has been shown that some PBP can interact with the WTA [9], again suggesting some cross-talk between the WTA and the mechanism of action of the beta-lactam antibiotics.

WTA itself is a very interesting target for the development of new antibiotic candidates. Tarocins and tunicamycin, for example, were shown to act on the synthesis of the WTA by inhibiting the first enzyme of the biosynthesis pathways, TarO [10], with bactericidal effects on resistant bacteria and also with synergistic effects with beta-lactam antibiotics. Tunicamycin, not only inhibits TarO but also inhibits MnaA, a UDP-N-acetylglucosamine 2-epimerase that provides the appropriate concentrations of UDP-GlcNAc and UDP-ManNAc for the further steps of the biosynthetic pathway [11,12].

As highlighted by Walsh and Wencewicz, the antibiotics currently used are limited to only five major validated targets/pathways [13]. So, given the scarcity of targets and the relevance of the cell wall decoration for *S. aureus*, we report here the structure-based screening for new MnaA binders using a catalog of existing drugs in a repurposing strategy [14]. After molecular docking and careful visual inspection, we identified fifteen ligand candidates. Further experimental evaluation with quantitative DSF (qDSF) revealed ticarcillin, a β-lactam, as a weak binder for *S. aureus* MnaA (SaMnaA). An equilibrium molecular dynamics simulation confirmed a stable binding pose over 600 ns with strong interactions between ticarcillin carboxylate groups and SaMnaA R206 and R207 residues.

## 2. Materials and Methods

### 2.1 Molecular Docking and Ligand Selection

The molecular model used for docking calculations was based on the crystal structure for *S. aureus* MnaA, PDB Id 5ENZ [12], with the same structure preparation procedure as previously described [15]. Briefly, the server H^++^ [16] was used to define the protonation states of residues, and hydrogen atoms were added accordingly. The atom types and charges were defined according to AMBER FF14SB [17]. In this model, a UDP molecule was considered as a part of the ‘receptor’. The atomic charges were computed using an RESP fitting procedure after initial calculations using Hartree-Fock method with 6-31G* basis, as available in Gaussian 09 [18]. The substrate UDP-GlcNAc was prepared using the same method and considered as the ‘reference ligand’ during the docking calculations. The ZINC Drug Database (ZDD) was used for the screening. The MOL2 files were used as provided by the database using the default ZINC methods for ligand preparation [19]. The docking calculations were set up using `LiBELa`, a ligand- and receptor-based docking software developed in our group [20], using a pre-computed `DelPhi` [21,22] electrostatic potential for modeling the polar interactions and a 6-12 Lennard Jones potential to model van der Waals interactions [23]. The docking results were analyzed using the ViewDock tool available in UCSF Chimera [24] and promising candidates were selected based on the visual inspection of favorable interactions involving the ligand and the enzyme.

### 2.2 SaMnaA expression and purification

*S. aureus* MnaA was cloned from S. aureus SA16 gDNA sample kindly provided by Prof. Dr. Ilana Camargo. The SaMnaA gene was amplified using the primers forward 5’- CAGGGCGCCATGATGAAAAAGATTATGACCATATTTG-3’ and reverse 5’-GACCCGACGCGGTTATCAGAAATCGCTTGGTTTT-3’. The amplified gene was inserted in the pET-Trx1A vector using a strategy described elsewhere [25]. *E. coli* Rosetta (DE3) cells were transformed with the resulting plasmid and used for protein expression. A starting culture was grown overnight in LB medium supplemented with containing 50 μg/ml kanamycin. Afterward, the starting culture was used to inoculate LB medium containing 50 μg/ml kanamycin. The cells were incubated in a shaker at 18°C and, after reaching an Optical density of 0.8 at 600 nm, the protein expression was induced with 0.1 mM IPTG. After 16 hours at 18°C, the cells were harvested by centrifugation at 7800 x g for 30 min and disrupted by sonication in lysis buffer containing 50 mM NaH_2_PO_4_, pH 8.0, 300 mM NaCl, 10 mM imidazole, 1 mM PMSF, 1 mg/mL lysozyme. The soluble proteins were isolated from the cell debris by centrifugation at 18,000 x g for 30 min at 4°C and SaMnaA was purified by IMAC. Briefly, the soluble fraction was incubated with 5 ml of the Ni resin equilibrated with washing buffer containing 50 mM NaH_2_PO_4_, pH 8.0, 300 mM NaCl, 20 mM imidazole, 10% glycerol. After extensive washing, MnaA was eluted in the same buffer supplemented with 500 mM imidazole. After dialysis to remove the imidazole, and concentration up to about 2 mg/ml and incubated with a Tobacco Etch Virus (TEV) protease for 16 hours at 4°C, to remove the fusion protein. A second IMAC step was carried out to remove the His-tagged fusion protein thioredoxin. For crystallization trials, an additional purification step was carried out with size exclusion chromatography using a Superdex 75 column.

### 2.3 Differential Scanning Fluorimetry (DSF)

Fourteen ligands were initially selected from docking calculations and were screened for their ability to bind to *S. aureus* MnaA *in vitro* using a DSF assay. Briefly, SaMnaA at 2μM was incubated with each ligand at a concentration of 6 μM (1:3 ratio) for 30 minutes. Afterward, the complex was incubated with SYPRO Orange dye (ThermoFisher) for DSF experiment using a Bio-Rad CFX96 real-time PCR Detection System equipment using a temperature range of 25-95 °C with a heating rate of 1°C every 30 seconds. The obtained data were adjusted according to the unfolding model proposed by Bai and coworkers [26]. Melting temperatures (T_m_) were extracted from an initial fitting to the DSF data, using ΔC_P_ as constant at 3.22 kcal/(mol.K), as estimated by the ScooP server [27], and used to evaluate actual binders among the ligands selected from docking calculations.

For affinity constant calculations, the same protocol was used, varying the ligand concentration from 0 to 2,000 μM. The thermal unfolding data were adjusted in the model proposed by Bai and coworkers [26]. In this model, the fraction of unfolded protein *f*_*U*_ is related to the ligand affinity Kd as:

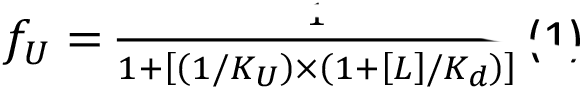

where [L] and KU are the free ligand concentration and the unfolded/folded equilibrium constant, respectively. After redefining the free ligand concentration [L] as a function of the total ligand concentration [L]_T_, *f*_*U*_ can be analyzed in terms of only two parameters, K_U_ and K_d_, since [L]_T_ and [P] are known [26]. All the data analysis was done using `DSF_fitting.py` python script as described by Bai and coworkers [26]. All the tested compounds were purchased from Merck-Sigma.

#### Molecular Dynamics Simulations

The docking pose found for Ticarcillin bound to *S. aureus* MnaA was used for equilibrium MD simulations. Here, the same receptor preparation described was used. For the ligand, RESP charges were computed after electrostatic potential calculation using HF/6-31G*, using the same protocol as used for UDP and UDP-GlcNAc. Ligand atom types were attributed according to GAFF2 [28]. The complex, containing MnaA and ticarcillin was immersed in OPC solvation box with 150 mM NaCl and neutralized using `tLeap`. The system was used for energy minimization, heating, and equilibration using the same protocol as previously described [15]. After, 5,000 minimization steps, the system was heated to 300 K in 50 ps using an NVT ensemble simulation, followed by 50 ps of an NPT simulation to equilibrate the system density. For these preliminary equilibration steps, harmonic restraints were kept in the initial coordinates of the ligands and the protein with a spring constant of 2.0 kcal/(mol.Å^2^). Finally, 500 ps of NPT simulation was carried out for final equilibration, followed by a 600 ns productive simulation (NPT). For the final equilibration and productive simulation, a weak positional restraint was kept at all α carbon atoms to avoid enzyme opening with a spring constant of 1.0 kcal/(mol.Å^2^). All the MD simulations were run with SANDER, PMEMD, and PMEMD.CUDA programs from the AMBER 20 package [29–32]. The MD trajectory was analyzed using AMBERENERGY++ [33], a locally written program to compute interaction energies, VMD [34], Chimera [24], and CPPTRAJ [35].

## Results

Given the relevance of *MnaA* in the synthesis of the WTA and the antibiotic relevance, we used a structure-guided screening of compounds that could inhibit the enzymatic activity. For this purpose, we used the ZINC Drug Database (ZDD), a catalog of almost 3,000 purchasable drugs that were approved for use in humans in different countries [19]. The rationale here involved the identification of binders that are already proved to be non-toxic and with a tolerable pharmacokinetic profile. In a typical drug discovery pipeline, the compounds could be further improved to improve their properties, including the affinity for the target. In this context, the identification of a binder in the narrow chemical space would be a challenge *per se*.

In our previous MD-based analysis of the domain movement in *S. aureus* MnaA, we found that the interaction between the enzyme and the substrate UDP-GlcNac is highly dependent on electrostatic interactions involving the residues R206, R207, K239, H40, and H238 within the active site [15]. Since the docking screen aimed to find new inhibitors, our inspection of the docking compounds prioritized molecules with polar groups (negatively charged or hydrogen bond acceptors) interacting with those residues. Also, compounds with Hodgkin’s similarity index smaller than 0.5 were deprioritized, as a usual indicator of an incomplete filling of the binding pocket. Finally, all the compounds with binding energies ranging from −50 to −20 kcal/mol were visually inspected in UCSF Chimera, following the prioritization rules described above.

After visual inspection of the docking results, the following compounds were selected for experimental evaluation: methocarbamol, pantoprazole, 5-azacytidine, pyridoxal 5’-phosphate, sulfamethizole vetranal, suprofen, sitaxentan, cytarabine, (e)-5,5’-(diazene-1,2-diyl)bi, ticarcillin, nilutamide, adenosine, thymidine, tolbutamide, L-Histidine, sulfadiazin vetranal, and sulfamethoxazole. The selection criteria involved the number of favorable interactions with at least two for the polar residues in the MnaA active site (R206, R207, K239, H40, and H238), the absence of ligand polar groups lacking interactions with the receptor, and a reasonable match in the occupied volume as compared to the substrate pose.

### Experimental Screening of Ligand Candidates

A rapid assay for screening of the ligand candidates was set up using a DSF assay. Initially, the enzyme melting curve was characterized in the apo state (unbound) as well as bound to UDP, UDP-N-Acetylglucosamine, and UDP + UDP-N-acetylglucosamine together. As indicated in Figure 1, the melting temperatures (T_M_) for the apo MnaA and the enzyme bound to UDP do not change significantly (54.4 and 55.2°C, respectively), suggesting that (i) the enzyme binds very weakly to UDP in the absence of the substrate or (ii) the enzyme could be pre-loaded with UDP after bacterial expression [12]. When bound to UDP-GlcNAc, on the other hand, an important shift in the melting temperature was observed, as shown in Figure 1, with a T_M_ of 58.2°C. Interestingly, when both UDP and UDP-N-acetylglucosamine were incubated with MnaA, the observed T_m_ was 58.7°C, very close to the T_m_ observed for UDP-N-acetylglucosamine alone.

**Figure 1.**
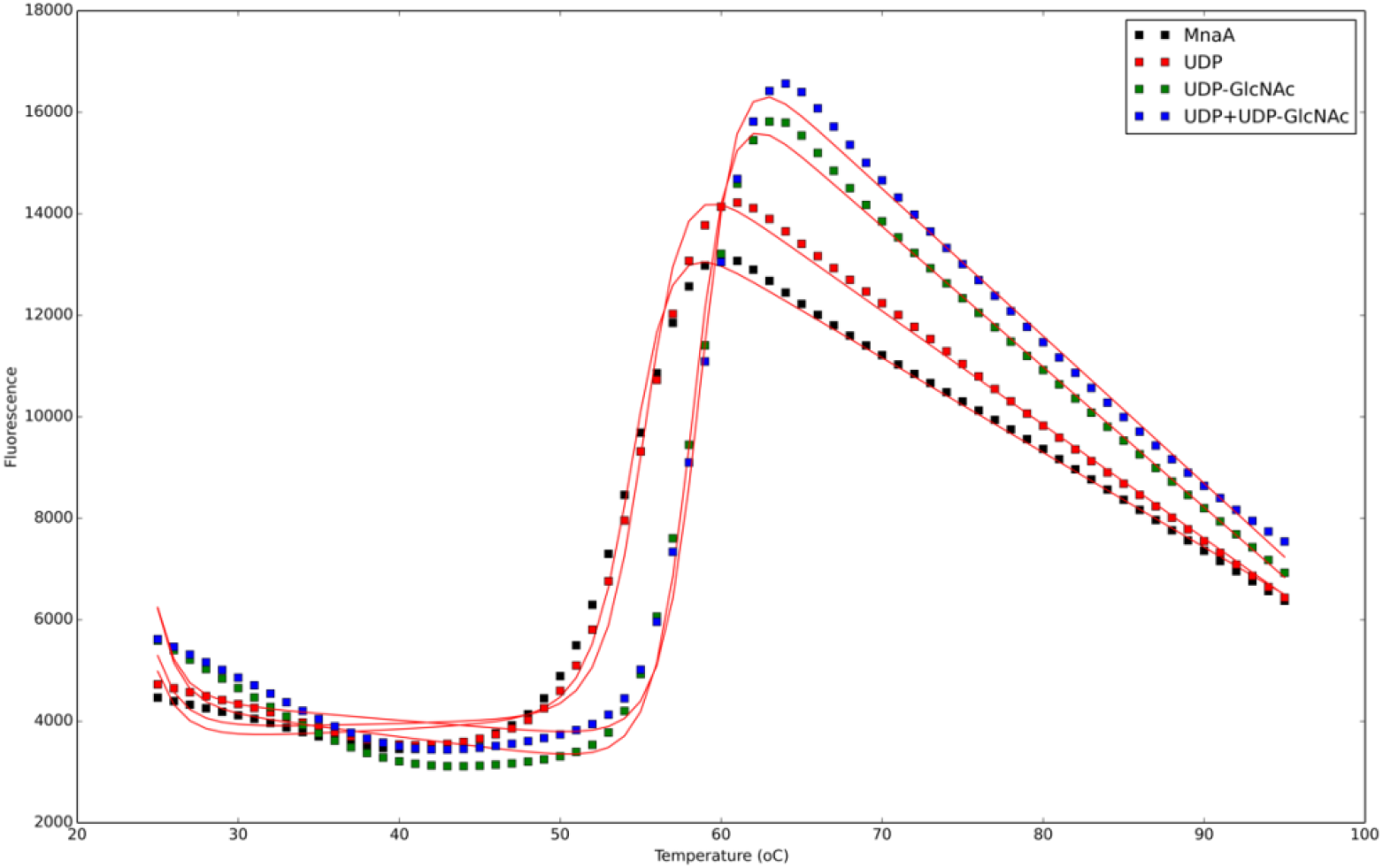
Melting curves for unbound (apo) MnaA (black squares) and in complex with UDP (red squares), UDP-GlcNAc (green squares), and UDP and UDP-GlcNAc simultaneously (blue squares). The red lines indicate the fit of the experimental data.

**Figure 2.**
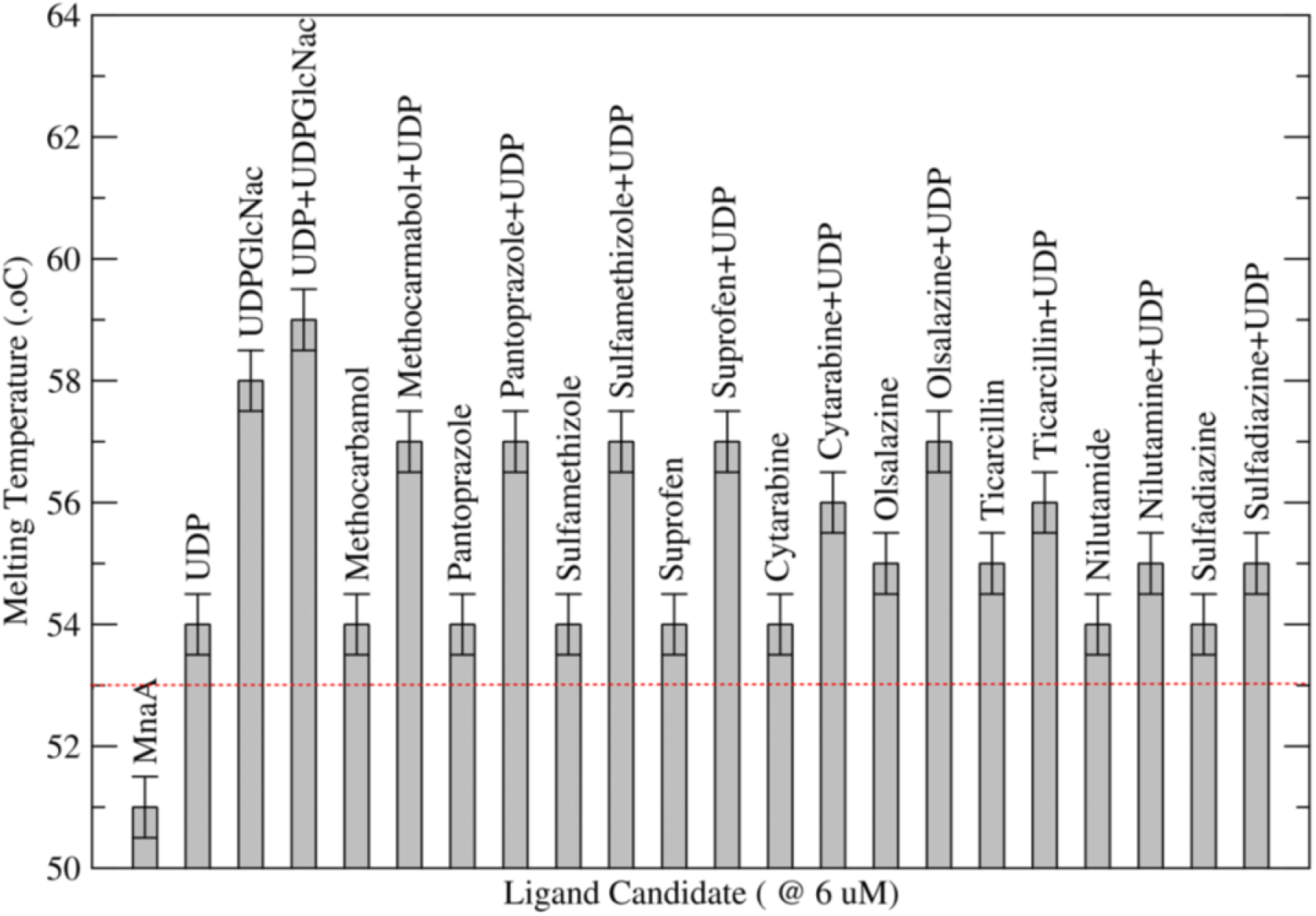
Initial hits in the experimental ligand screening. SaMnaA (2μM) and the ligands identified in the virtual screening (6μM) were preliminarily tested for their binding properties using the DSF assay. A thermal shift of 2 degrees (red line) was considered an indication of binding. In all cases, the incubation of the tested ligand with UDP increased the observed T_M_.

This preliminary result indicated that even the substrate, binds with a relatively small change in the T_m_, of about 4 degrees. So, to identify potential MnaA binders, we searched for ligands that resulted in a T_m_ change of at least 2 degrees in a DSF screening experiment. From the 17 initially purchased compounds, nine showed a T_M_ shift of 2 degrees or higher and were selected for further experimental evaluation. The ligand candidates initially tagged as potential hits were methocarbamol, pantoprazole, sulfamethiazole, suprofen, cytarabine, olsalazine, ticarcillin, nilutamide, and sulfadiazine, while sulfamethoxazole, tolbutamide, histidine, thymidine, pyridoxal 5’-phosphate were discarded as potential ligands. 5-azacytidine revealed fluorescence signal in the absence of protein and was removed from our analysis.

For further exploration of the binding potential of the selected compounds, a quantitative DSF assay was set up, as previously described by Bai and coworkers [26], where the change in T_M_ is monitored as a function of the ligand concentration. First, a positive control experiment was set up, using MnaA substrate, UDP-GlcNac. The analysis of the melting curves over a concentration range of 0 - 2,000 μM revealed a clear right-shift in the melting temperature (T_m_) for an increasing ligand concentration, as shown in Figure 3. According to the global fitting of the data, measured in two independent replicates, at T_m_=54°C, a K_U_ of 1.08 was achieved with a K_D_ of 33 μM. At T_m_=56°C, a K_U_ of 3.22 with a K_D_ of 31 μM was observed. Here, K_U_ is an equilibrium constant between the unfolded and folded enzyme. So, a K_U_ of 1 gives equal concentrations of folded and unfolded species. Interestingly, the K_M_ for UDP-GlcNAc measured for SaMnaA was reported to be 411 ± 57μM according to Mann and coworkers [12]. The differences can be due to the assumptions of an equilibrium interaction rather than a kinetic (enzymatic) model. In line with this observation, a plot of ln *K* as a function of 1/*T* does not show a typical linear dependency expected for the van’t Hoff relationship, indicating that the enthalpy may not be temperature independent. This could be related to the change in enzyme activity over the temperature range, resulting in this non-linear relationship.

**Figure 3.**
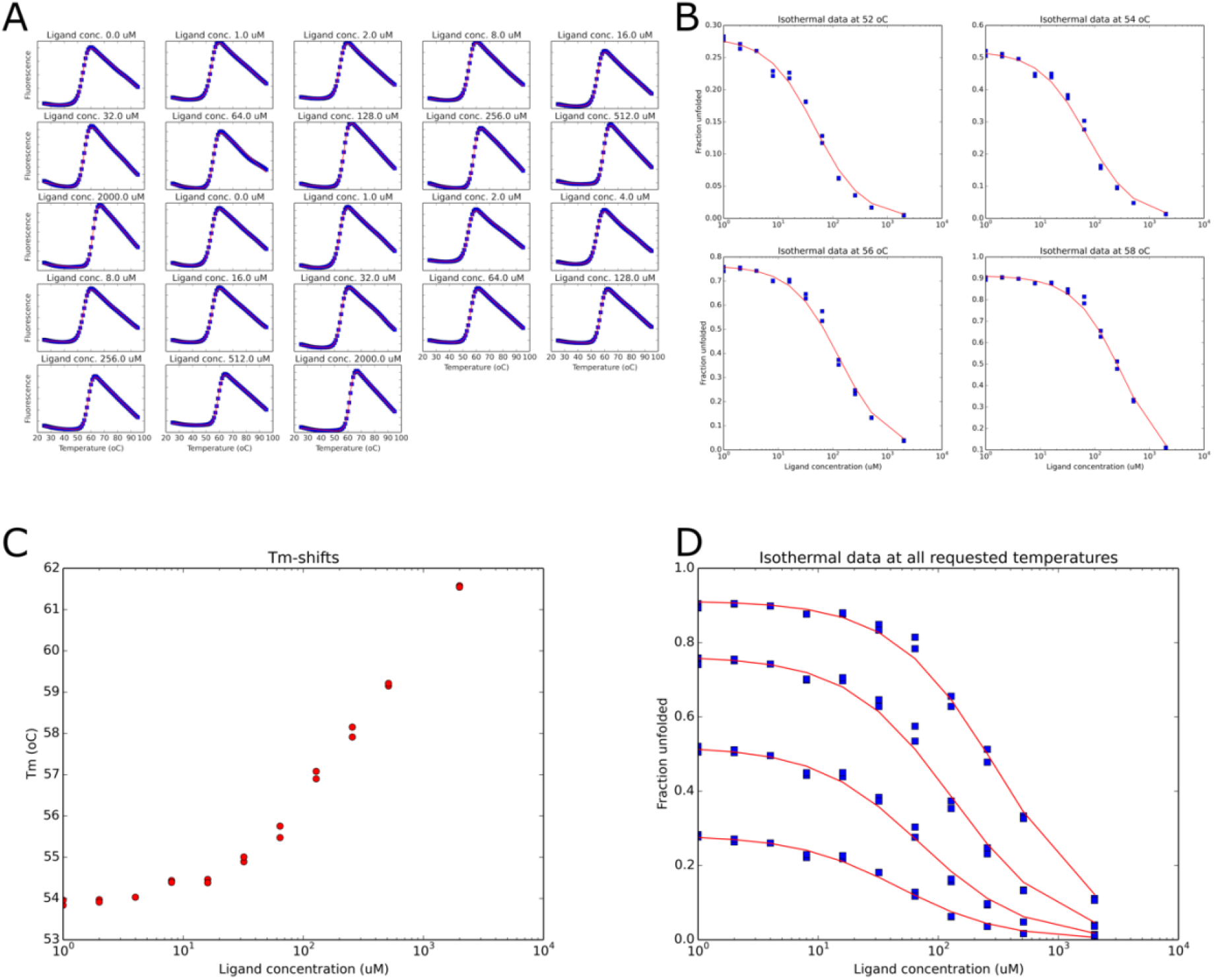
Isothermal analysis of UDP-GlcNAc binding to SaMnaA based on DSF data. (A) Thermal denaturation fluorescence data and fitting curve (red lines) for different ligand concentrations. (B) Fraction of unfolded protein as a function of UDP-GlcNac at different temperatures. (C) Shifts observed in the melting temperature T_M_ as a function of ligand concentration. (D) Overlay of the data shown in panel (B) of the fraction of unfolded protein as a function of ligand concentration. In all cases, the red lines show the best fit obtained from the models shown in equation (1).

For most of the ligands selected in the initial screen, a clear T_M_ shift as a function of the ligand concentration could not be observed. Instead, a constant T_M_ was observed as a function of the ligand concentration, with a sharp increase in fluorescence in higher concentrations. This sharp increase in fluorescence can indicate some interaction between the ligand and fluorescent probe, for example.

For ticarcillin, however, the melting curves were measured in two replicates over a concentration range of 0-1,300 μM, limited by the ligand solubility, and at T_M_=54°C a K_U_ of 1.08 was observed with a K_D_ of 421 μM (Figure 4). For ticarcillin, in contrast to what was observed for UDP-GlcNAc, a linear relationship was observed for ln *K* as a function of 1/*T*.

**Figure 4.**
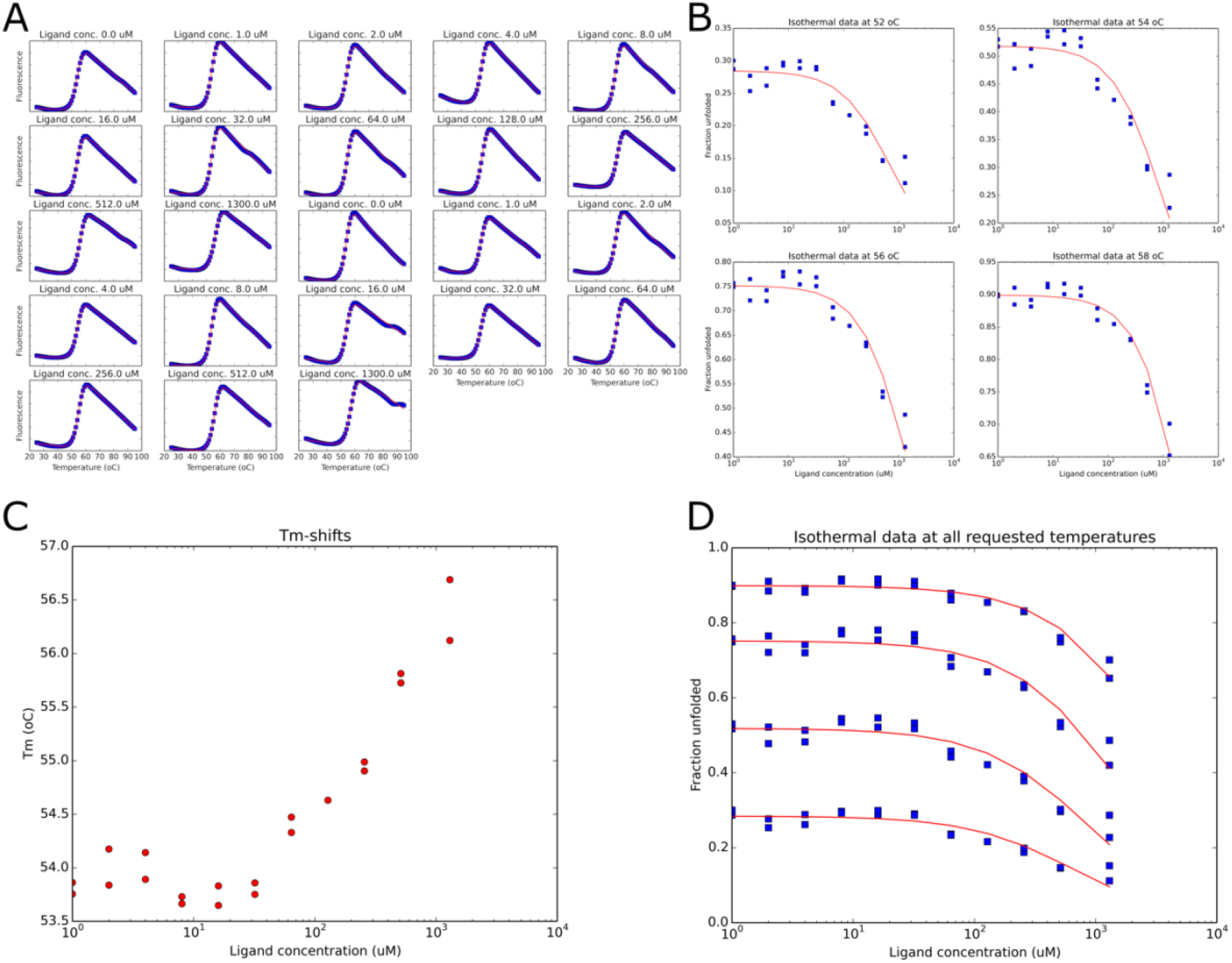
Isothermal analysis of ticarcillin binding to SaMnaA based on DSF data. (A) Thermal denaturation fluorescence data and fitting curve (red lines) for different ligand concentrations. (B) Fraction of unfolded protein as a function of ticarcillin at different temperatures. (C) Shifts observed in the melting temperature T_M_ as a function of ligand concentration. (D) Overlay of the data shown in panel (B) of the fraction of unfolded protein as a function of ligand concentration. In all cases, the red lines show the best fit obtained from the models shown in equation (1).

The quantitative DSF data indicates that ticarcillin is a weak MnaA binder with a K_D_ in the range of 400 μM, in our experimental conditions, which is about thirteen times greater than the K_D_ measured for the substrate UDP-GlcNAc in our conditions. Although the quantitative DSF approach does not measure the binding constant in an equilibrium situation, previous data have shown that binding affinities agree very closely to the binding affinities determined by other experimental approaches, such as Isothermal titration calorimetry (ITC) [26].

### Molecular Dynamics Simulation of the SaMnaA-Ticarcillin Complex

The docked pose from ticarcillin bound to SaMnaA was used to generate the initial coordinates of the solvated complex for an equilibrium MD simulation. The obtained 600 ns trajectory was used for cluster analysis in CPPTRAJ [35] and also in UCSF Chimera. In both cases, two major clusters were identified with minor differences among them. Figure 5 shows the ligand bound to the SaMnaA, as observed in the most populated cluster in the MD simulation.

**Figure 5.**
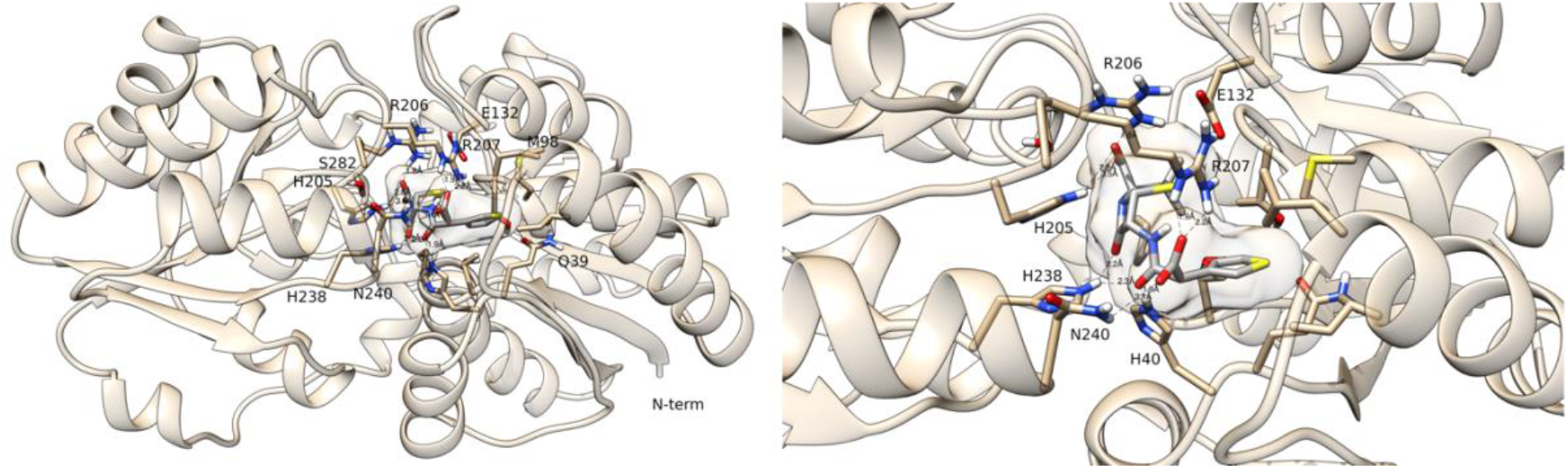
A representative frame of the most populated cluster during the MD simulation of the SaMnaA-ticarcillin complex. Solvent and non-polar hydrogen atoms were omitted for clarity. Ticarcillin is shown in gray sticks.

As shown in Figure 5, several polar residues interact with ticarcillin in the binding site, contributing to its binding affinity. Most importantly, H40, N240, H238, H205, R207, and R206 directly Interact with ticarcillin carboxylate groups as well as with carbonyl oxygen. As previously described, it is interesting to note that many positively charged residues surround the binding site, which would be expected for an enzyme that binds phosphorylated substrates [15]. That said, ticarcillin seems to be a reasonable binder, given its total negative charge.

The individual contributions of each polar residue in the vicinity of the catalytic cleft were computed with `AmberEnergy++` and are shown in Figure 6 below. As evidenced in the left panel, the charged arginine residues have the largest contribution to the binding energy, with about −130 kcal/mol per residue, while the entire interaction energy between ticarcillin and the environment (MnaA and explicit solvent) sums about −450 kcal/mol (Figure 6, right panel). This finding is also in line with our previous analysis that showed that R206, R207, and K239 were the residues with the largest contribution to UDP-GlcNac binding to MnaA [15]. Here, K239 contributes about −50 kcal/mol to the ticarcillin binding energy.

**Figure 6.**
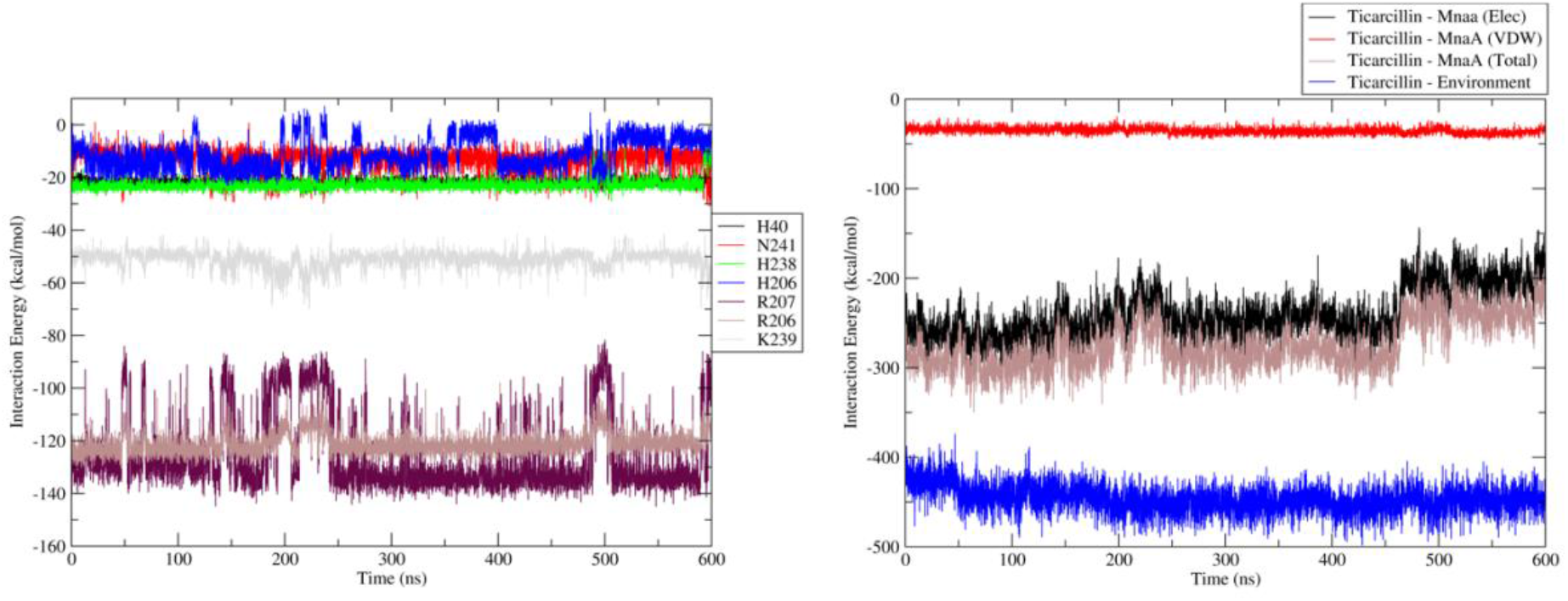
Interaction energies among ticarcillin and MnaA. (left) Decomposition analysis of the individual contributions to the binding energy for the polar residues in the vicinity of the active site. (right) Interaction energies computed for ticarcillin and the entire protein, as well as its electrostatic and van der Waals terms. The blue line shows the interaction energy of the ligand (ticarcillin) with the environment, i.e., protein and the explicit solvent.

Additional analysis of the interaction energy with MMPBSA.py [36] revealed an interaction energy of −47 ± 6 kcal/mol, using a Generalized Born (GB) model from Hawkins, Cramer, and Truhlar [37]. In line with the previous analysis, ticarcillin binding to SaMnaA has a van der Waals contribution of −35 ± 4 kcal/mol, while the electrostatic contributions account for −239 ± 28 kcal/mol. The GB correction for the (implicit) solvent adds +232 ± 24 kcal/mol, resulting in a final binding energy of −47 kcal/mol.

## Discussion

The limited number of molecular targets/pathways explored by existing antibiotics is worrying [13]. The resistance problem is even more severe for some pathogens where high resistance rates are observed. The data reported to the National Healthcare Safety Network 2015-2017 indicated, for example, that 48.4% of the *S. aureus* isolates from device-associated healthcare associate infections in acute-care hospitals were found to be resistant to methicillin (MRSA) [38]. 82.1% of the *Enterococcus faecium* isolates were found to be vancomycin-resistant and 43% of *Acinetobacter spp* were found to be multidrug-resistant [38]. These alarming trends highlight the need for new antibiotic targets and the basic and translational research on their mechanisms.

Here we described the structure-based identification of ticarcillin as a weak MnaA binder. According to the docking model and the binding mode observed in the MD simulations, ticarcillin binds to the substrate pocket, putatively acting as a competitive inhibitor, although this proposal still awaits further experimental validation. This finding has at least two interesting implications. The first implication is the usage of ticarcillin as a scaffold for the development and optimization of compounds aiming for higher affinity to SaMnaA. Those compounds can be useful as chemical probes to better understand the effects of SaMnaA inhibition in *S. aureus* cultures. Since ticarcillin is a β-lactam antibiotic, there is no sense in testing it since a reduction in cell proliferation is already expected to be observed.

The second implication is associated with the ticarcillin mechanism of action. Interestingly, this β-lactam is expected to reach serum concentrations of up to 124 μg/ml after an intravenous injection of 1 g [39], which means serum levels even between 300 and 400 μM. So, at these concentrations, MnaA could be expected to be partially bound to ticarcillin. This off-target interaction for ticarcillin could explain, at least partially, the good activity of ticarcillin on the Gram-negative bacteria *Pseudomonas aeruginosa*. *P. aeruginosa* has a UDP-N-Acetylglucosamine 2-epimerase that shares about 51% in sequence identity with *S. aureus* MnaA. The same is true for species of the *Acinetobacter* genus. Thus, it is tempting to speculate that the MnaA inhibition may be associated with the ticarcillin activity on Gram-negative bacteria. In line with this observation, it is interesting to note that the activity on Gram-negative bacteria is only achieved at high concentrations [39], again suggesting a low-affinity interaction with the target.

If this hypothesis is correct, is it possible to anticipate other pathogens that could be susceptible to ticarcillin? According to a sequence comparison in BLAST [40], other UDP-N-acetylglucosamine 2-epimerases are close orthologues of *S. aureus* MnaA, including a *Mycoplasma* epimerase. Since *Mycoplasma* species are usually unresponsive to beta-lactams, it may be a feasible experimental model to assess whether the UDP-N-acetylglucosamine can inhibit the growth and, maybe, indicate a new therapeutic indication for ticarcillin. Although this experiment is not accessible to us, we foresee very promising consequences of the findings reported here.

## Funding

We thank Fundação de Amparo à Pesquisa do Estado de São Paulo – FAPESP, for the financial support via grants 2020/03983-9, 2017/18173-0, 2015/26722-8, and 2015/13684-0; and the Conselho Nacional de Desenvolvimento Científico e Tecnológico – CNPq via grant 303165/2018-9. This study was also financed in part by the Coordenação de Aperfeiçoamento de Pessoal de Nível Superior – Brazil (CAPES) – Finance Code 001.

## Declaration of Competing Interests

The authors declare that they have no known competing financial interests regarding the work described in this paper.

## Acknowledgments

The authors thank Maria A. M. Santos, Lívia R. M. Margarido, Josimar Sartori, and João F. Possatto, for their technical assistance. We are also indebted with Luana G. Morão for her critical reading of this manuscript. We thank prof. Ilana Camargo for kindly providing the genomic DNA for *S. aureus* SA 16.

## Author contributions

ASN wrote the original draft and supervised and administrated the project. ASN and ECA were involved in the hypothesis generation, definition of the methodology, data acquisition, and analysis. The final manuscript was revised by ASN and ECA.

## Data accessibility

The data generated in this study are available from our group’s GitHub account at https://github.com/alessandronascimento/LabData/tree/main/MnaA-Ticarcillin.

## Notes

https://github.com/alessandronascimento/LabData/tree/main/MnaA-Ticarcillin

### Competing Interest Statement

The authors have declared no competing interest.

### Summary of Updates

Some additional details about the ligand selection were added and small typos were corrected.

